# Mobile phones: The effect of its presence on learning and memory

**DOI:** 10.1101/678094

**Authors:** Clarissa T. Tanil, Min Hooi Yong

**Affiliations:** Department of Psychology, Sunway City, Selangor, Malaysia

## Abstract

Our aim was to examine the effect of mobile phone’s presence on learning and memory among undergraduates. A total of 119 undergraduates completed a memory task and the Smartphone Addiction Scale (SAS). As predicted, those without phones had higher recall accuracy compared to those with phones. Results showed a significant negative relationship between phone conscious thought and memory recall but not for SAS and memory recall. Phone conscious thought significantly predicted memory accuracy. We found that the presence of a phone and high phone conscious thought affects one’s memory learning and recall, indicating the negative effect of a mobile phone proximity to our learning and memory.

## Introduction

Today, mobile devices such as smartphones are a popular communication medium and is likely to be the most dominant form of communication, especially among adolescents. Smartphone ownership in adolescents and young adults is 95% in the United States, 46% in Western Europe, 72% in South Korea, and 71% in Taiwan (1,2). The phone has evolved from basic communicative functions e.g. calls only, to being almost a computer-replacement device used for web browsing, instant communication on social media platforms and productivity tools, e.g. word processing etc. Undoubtedly, the constant connectivity is applauded and desired but this has also spiralled into an obsession with the device for many individuals (3). The consequence of high smartphone usage is a current source of debate, considering its potential to be a distractor in one’s life.

### Mobile phone as a distractor

Altmann, Trafton, and Hambrick (4) suggested that as little as a 3-second distraction (i.e. reaching for a phone) is adequate to disrupt attention while performing a cognitive task. The distraction caused by the phone is disadvantageous to subsequent tasks; creating more error as the distraction period increases, and is particularly evident in classroom settings. While teachers and parents are for (5) or against phones in classrooms (6), empirical evidence showed that students who used their phones in class took fewer notes (7), and had overall poorer academic performance compared to those who did not (8,9). Bowman, Levine, Waite and Gendron (10) reported no significant difference in-class test scores that measured comprehension between those who texted in class and those who did not. Yet, texters took a significantly longer time to complete the in-class test compared to those who did not, suggesting that texting students were simply too focussed on texting, hence requiring more cognitive effort in memory recall.

### Mobile phone presence and working memory

Recent studies demonstrated that the mere presence of a mobile phone may have detrimental effects on cognitive functioning such as learning and memory. Thornton, Faires, Robbins and Rollins (11) tested undergraduates on digit cancelation tasks (simple and demanding versions) which measured attention and cognitive capacity and a trail making test to measure attentional processes. A questionnaire was also administered to measure mobile phone usage. Students were divided into two groups; a mobile phone or a phone-sized notebook placed on participant’s table before completing the tasks. Results showed no significance on performance between the phone and notebook condition during simple version of the digit cancellation task. However, participants with the mobile phone group had poorer performance on the demanding version of the digit cancellation task compared to those with the notebook. In another study, the researchers tested students in three experimental conditions: phones located within reach (high salience/HS), phone located in a different room from the participant (low salience/LS) and a control group (the phones were kept in the bags or pockets) (12). The students completed the Automated Operation Span (OSpan) task and Raven’s Standard Progressive Matrices to measure their cognitive capacity and fluid intelligence. They were also asked to complete a questionnaire to determine the degree of phone usage. Results showed that LS participants performed significantly better in the tests compared to HS condition while no difference was found between the control condition and HS or LS conditions. Their findings led to Ward et al. concluding that having a phone within one’s sight increases cognitive load because greater cognitive effort is required to inhibit distractions posed by the phone. Unlike Thornton’s study in which there were no mediating or moderating effects in students’ performance with phone usage, Ward et al. (12) reported that their participants’ performance in the tasks were moderated by the degree of attachment/dependence to their phones. Ward et al. also measured the participant’s perception on the degree to which their phone may have affected their performance. Results showed that most participants were unaware of the effect that their phone might have on performance in the memory task. In another study, participants were divided into two conditions; mobile phone or paper notebook hung on the left side of the computer screen (13). Participants completed a visual spatial search “T” either from an 8 or 24 stimuli display and two questionnaires; a mobile phone usage and attachment, and Internet addiction. Results showed that participants in the phone condition had slower reaction times compared to the notebook group in both 8 and 24 stimuli and that those who scored high in the mobile phone usage and attachment questionnaire could rapidly identify the target at the congruent location, but not for the low scores group. In sum, these findings suggest that the mobile phone has become ubiquitous in an individual’s life to the point that one tends to discount the effect of this device on their lifestyle, especially on attention and memory even when not in use.

### Mobile phone addiction and mood states

Reliance of mobile phones has been linked to a form of psychological dependency. Anxiety arising from separation from these devices can interfere with one’s ability to attend to information. Cheever, Rosen, Carrier, and Chavez (14) observed that college students are most susceptible to the undesirable effects of overuse. High mobile phone use was found to be linked with higher chance of experiencing phone separation anxiety or also known as “nomophobia or no mobile phone phobia”, an anxiety characterized by constantly thinking about their phones and the desire to stay in contact with it (15). One study reported that participants with high phone attachment experienced separation-anxiety, with an increased tendency to attend to separation-related stimuli e.g. cupboard that stored their phones (16). Participants experienced feeling more anxious and unpleasantness when prohibited from using the phone compared to those who could (17). This is concordant with another study when the participants had to give up their smartphone for a day (18). Together with Ward et al. (12) findings, phone usage and its proximity to us is strongly related to our mood, and the absence of a mobile phone seems to induce feelings of anxiety and emotions of negative valence.

Moreover, ones’ mood may indirectly influence cognitive functioning. Payne and Schnapp (2014) examined undergraduates’ mood with the Positive and Negative Affect Scale (PANAS) and Cognitive Failures Questionnaire. They found that higher negative mood increased cognitive failures such as memory and attention. But some argued that being in a positive mood state improves creativity and increases flexibility for task switching tasks (19). In another, participants in either happy or sad moods performed poorly in a memory recall task, compared to the neutral condition (20). Overall, these findings suggest that the complex relationship between mood states and attachment to phone together with high usage might be a mediator to one’s cognitive performance.

### Present Study

Our aim was to examine the effects of mobile phone presence on learning and memory in undergraduate students. We also investigated the relationship between mobile phone addiction, mood states and attachment to memory recall accuracy. We hypothesised that (H1) participants in the “phone absent” (LS) condition are more likely to have higher memory accuracy compared to those in the “phone present” (HS) condition, (H2) participants with higher mobile phone addiction score and higher phone conscious thought are more likely to have lower memory accuracy, and (H3) LS participants will report an increase of negative affect or a decrease in positive affect after experiencing separation from their phone. Furthermore, (H4) we examined how these predictor variables; mobile phone addiction, phone conscious thought and affect/mood differences predict memory accuracy.

## Materials and Methods

### Participants

A total of 119 undergraduate students (61 females, *M*_*age*_ = 20.67 years, *SD*_*age*_ = 2.44) were recruited from a private university in an Asian capital city. To qualify for this study, the participant must own a smartphone, and must not have any visual or auditory deficiencies. Participants did not receive any compensation for participation. This study was approved by the Department of Psychology Ethics Committee (20171090).

Out of the 119 participants, 43.7% reported using their phone mostly for social networking, followed by communication (31.1%) and entertainment (17.6%) (see Table 1 for full details on phone usage). Participants reported an average phone use of 8.16 hours in a day (*SD* = 4.05). There was no significant difference between daily phone use for participants’ in the high salience (HS) and low salience groups (LS), *t* (117) = 1.42, *p* = .16. Female participants spent slightly more time using their phones over a 24-hour period (*M* = 9.02, *SD* = 4.10) compared to males, (*M* = 7.26, *SD* = 3.82), *t* (117) = 2.42, *p* = .02.

**Table 1.** Most frequently used phone feature (n=119).

|  | n | % |
| --- | --- | --- |
| Social networking (Instagram, Twitter, Facebook) | 52 | 43.7 |
| Communication (WhatsApp, Line, messaging, calls, emails) | 37 | 31.1 |
| Entertainment (music, games, videos) | 21 | 17.6 |
| Web surfing | 8 | 6.7 |
| Productivity (camera, calculator, alarm, calendar) | 1 | 0.8 |

### Study design

Our experimental study was a mixed design with phone presence (present, absent) as a between-subject, and memory task as a within-subject. Participants who had their phone out of sight from the participant ‘Absent’ or low-phone salience (LS), and the other group had their phone placed next to them throughout the study ‘Present’ or high-phone salience (HS). The dependent variable was recall accuracy from the memory test.

### Stimuli

#### Working memory span test

A computerized memory span task retrieved from software Wadsworth CogLab 2.0 was used to assess working memory (21). A working memory span test was chosen as a measure to test participants’ memory ability for two reasons. First, participants needed to learn and memorize three stimulus type which serves as the learning and memory component. Second, task completion takes approximately 20 minutes on average. This was advantageous because we wanted to increase separation-anxiety (16) as well as having the most pronounced effect on learning and memory without their phone’s presence (9).

The test comprised of three stimulus types such as words (long words such as computer, refrigerator and short words like pen, cup), letters (similar sound E, P, B and non-similar sound D, H, L) and digits (1 to 9). The test begun by showing a sequence of items on the left side of the screen, with each item presented for one second. After presentation, participants were required to recall the stimulus from a 9-button box located on the right side of the screen. In order to respond correctly, participants were required to click on the buttons for the items in the corresponding order they were presented. A correct response increases the length of stimulus presented by one item (for each stimulus category) while an incorrect response decreases the length of the stimulus by one item. Each trial began with five stimuli and increased or decreased depending on participants’ performance. The minimum length possible was one while the maximum was ten. Each test comprised of 25 trials with no time limit but without break between trials. Working memory ability is measured through the number of correct responses over total trials: scores range from 0 to 25, with the highest score representing superior working memory.

#### Positive and Negative Affect Scale (PANAS)

We use PANAS to assess the current mood of the participants (22). This questionnaire measures participants’ current positive affect/mood (PA) and negative affect (NA) state through feeling-describing statements. PANAS comprises of ten PA statements, such as “interested, enthusiastic, proud” as well as 10 NA statements, such as “guilty, nervous, hostile”. Each statement is measured though a 5-point Likert scale ranging from very slightly or not at all to extremely. Scoring is achieved by summing the ratings for each positive and negative affect which results in score ranging from 10 to 50, with higher scores representing higher levels of positive or negative affect. In the current study, the internal reliability of PANAS is good with a Cronbach’s alpha coefficient of .819, and .874 for PA and NA respectively.

#### Smartphone Addiction Scale (SAS)

SAS is a self-report scale used to examine participants’ smartphone addiction (23). SAS consists of six sub-factors whereby each factor is described through statements and scored through a six-point Likert scale ranging from strongly disagree to strongly agree. The sub factors include daily-life disturbance, positive anticipation, withdrawal, cyberspace-oriented relationship, overuse, as well as tolerance. “Daily-life disturbance” sub-factor measures the extent to which mobile phone use impairs one’s activities during everyday tasks, with sample questions such as “Missing planned work due to cell phone use” as well as “Feeling tired and lacking adequate sleep due to excessive cell phone use”. Secondly, “positive anticipation” is used to describe the excitement of using phone and de-stressing with the use of mobile phone. This is explored by asking questions such as “Feeling pleasant or excited while using a cell phone” or “Using a cell phone is the most fun thing to do”. “Withdrawal” sub-factor involves statements describing the feeling of anxiety when separated from one’s mobile phone. “Cyberspace-oriented relationship” sub-factors include questions pertaining to one’s opinion on online friendship. “Overuse” sub-factor raises statements designed to measure the excessive use of mobile phone to the extent that they have become inseparable from their device. Lastly, “Tolerance” was defined as the cognitive effort to control the usage of one’s smartphone. Scoring is achieved by summing all sub-factors in the scale. Scores range from 33 to 198. Higher SAS scores represent higher degrees of compulsive smartphone use. In the present study, the internal reliability of SAS was identified with Cronbach’s alpha correlation coefficient of .918.

### Procedure

We randomly assigned the participants one of two conditions: low phone salience (LS) and high phone salience (HS). Participants were tested in groups ranging between three to six people in a university computer laboratory, and each group was given the same experimental condition. They were seated two seats apart from each other to prevent communication. At the start of the experiment, participants were briefed on the rules in the experimental lab, such as no talking, and no phone use (for HS groups only). Participants were instructed to silence their phones. Each group was randomly assigned to either one of the experimental conditions, HS or LS. Participants in the HS condition were asked to place their smartphone on the left side of the table with the screen facing down. Meanwhile LS participants were asked to hand their phone to the researcher at the start of the study and the mobile phones were kept on the researcher’s table throughout the task at a distance between 50cm to 300cm from the participants depending on their seat location and out of sight behind a small panel on the table.

Participants first filled in the consent form and demographic form before completing a PANAS questionnaire. They were then directed to CogLab software and began the working memory test. Upon completion, participants were asked to complete the PANAS again followed by the SAS and about their conscious thought regarding their phone (“During the memory test how often do you think of your smartphone?”) on a scale of 1 (none to hardly) to 7 (all the time) and their perception on their phone use on their learning performance and attention span “In general, how much do you think your smartphone affects your learning performance and attention span?” on a scale of 1 (not at all) to 7 (very much). The researcher thanked the participants and returned the mobile phones in the LS condition at the end of the task.

## Results

### Phone presence and memory recall accuracy

An independent-sample *t*-test was used to examine whether participants’ performance on a working memory task was influenced by the presence (HS) or absence (LS) of their smartphone. Results showed that participants in the LS condition had higher accuracy (*M* = 14.21, *SD* = 2.61) compared to HS (*M* = 13.08, *SD* = 2.53), *t* (117) = 2.38, *p* = .02 (see Fig 1). The effect size ŋ^2^ = .44 indicates that mobile phone presence/salience has a moderate effect on participant working memory ability, and a sensitivity power of .66. We also examined gender difference on memory accuracy and found no significant difference between the males and females, *t* (117) = .18, *p* = .86.

**Fig 1.**
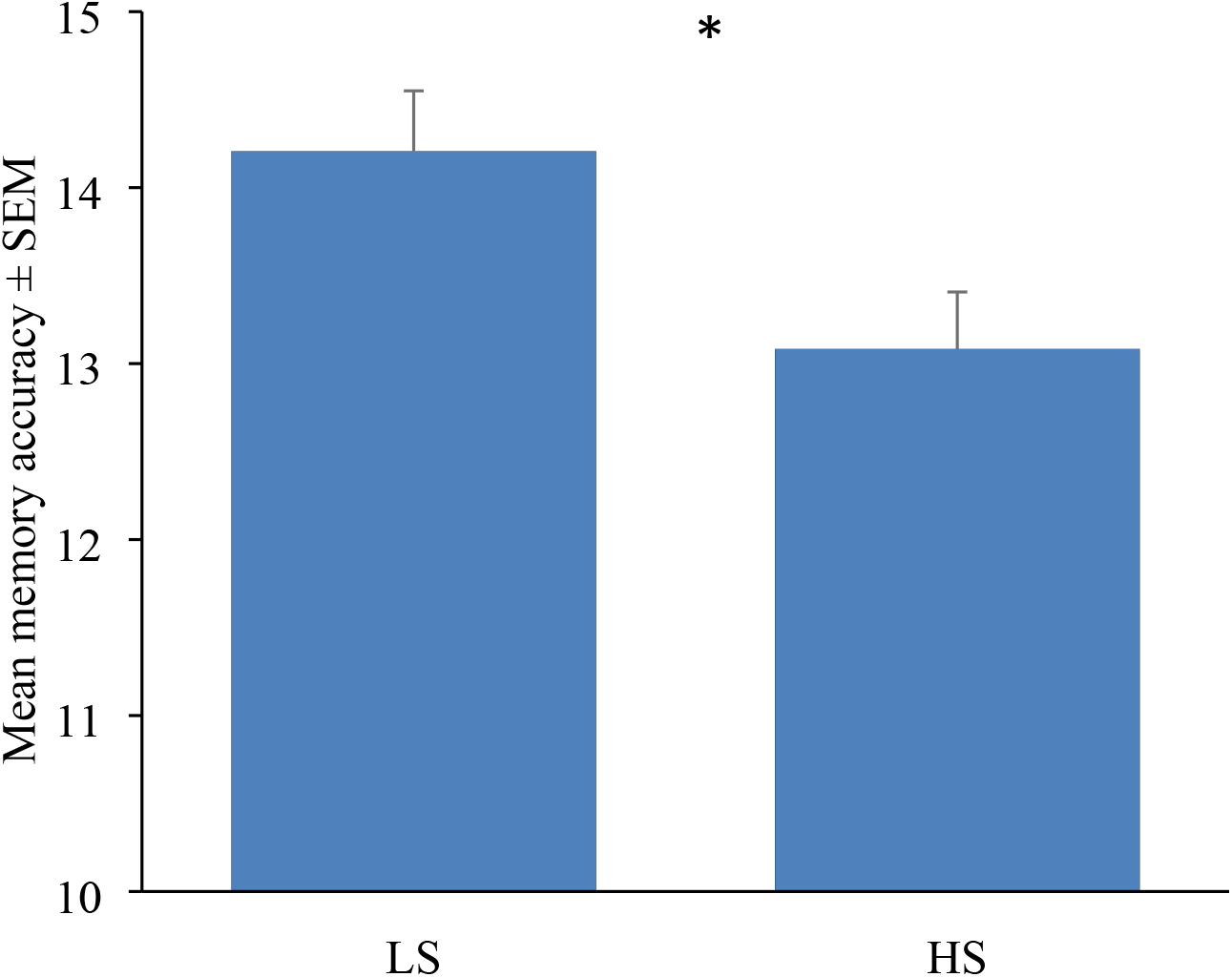
Mean memory accuracy between low phone salience (LS) and high phone salience (HS) groups (n = 119) * p < .05.

### Relationship between Smartphone Addiction Score (SAS) and higher phone conscious thought to memory recall accuracy

Participants reported an average of 103.65 (SD = 22.63) in the Smartphone Addiction Scale (SAS) (Kwon et al, 2013). There was no significant difference between the LS (*M* = 104.64, *SD* = 24.86) and HS (*M* = 102.70, *SD* = 20.45) SAS scores, *t* (117) = .46, *p* = .64 (see Fig 2).

**Fig 2.**
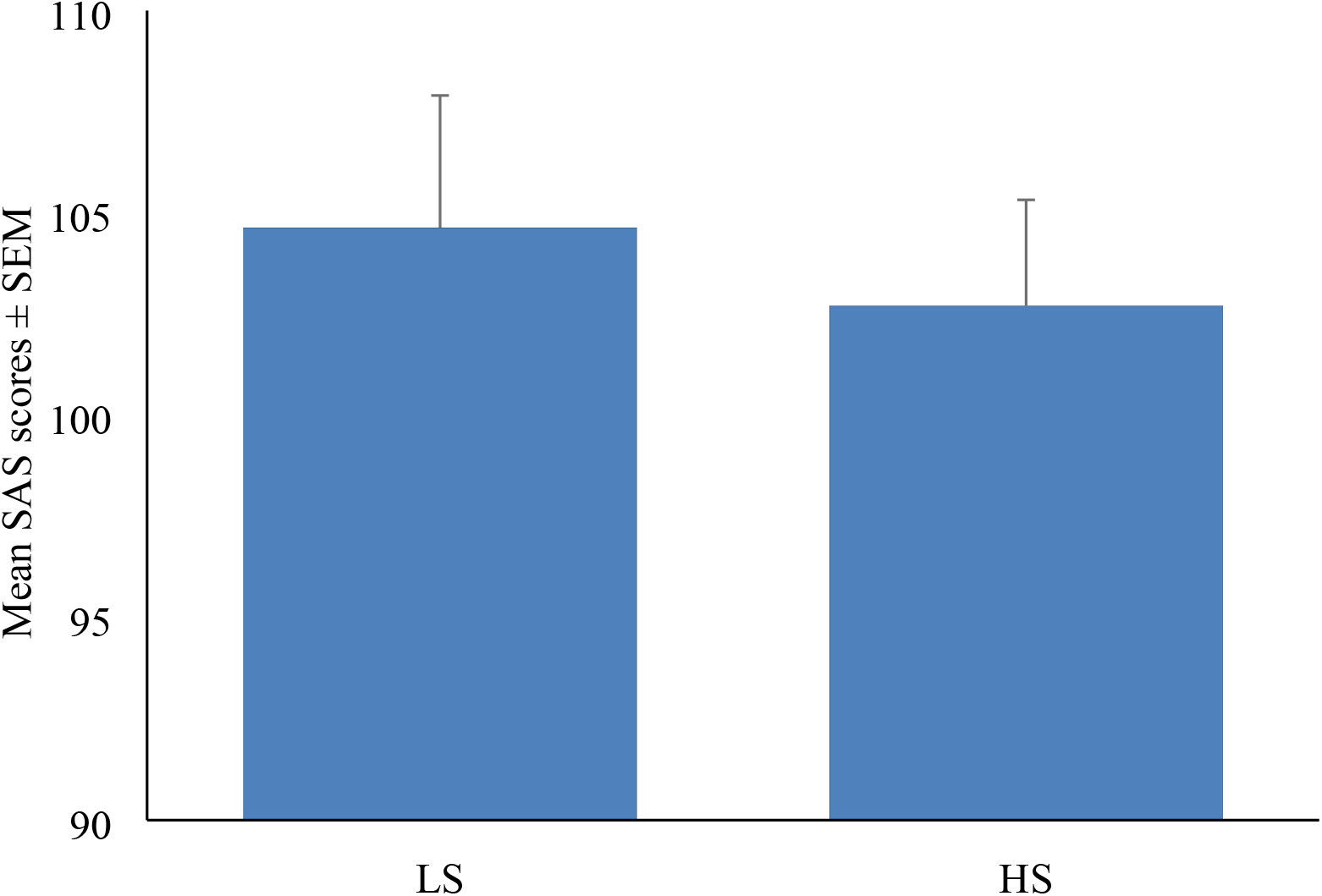
Mean Smartphone Addiction Scale (SAS) scores between low phone salience (LS) and high phone salience (HS) groups.

We predicted that those with higher SAS scores will have lower memory accuracy, and thus we examined the relationship between smartphone addiction (SAS) and memory accuracy using Pearson correlation coefficient. Results showed that there was no significant relationship between SAS and memory accuracy, *r* = −.03, *n* = 119, *p* = .76. We also examined the SAS scores between the LS and HS groups on memory scores. In the LS group, no significant relationship was established between SAS score and memory accuracy, *r* = −.04, *n* = 58, *p* = .74. Similarly, there was no significant relationship between SAS score and memory accuracy in the HS group, *r* = .10, *n* = 61, *p* = .47. We also found no significant relationship between each sub-factor of SAS scores and memory accuracy, all *p*s > .12 (see Table 2).

**Table 2.** Subscales of the Smartphone Addiction Scales (SAS) (n = 119)

| Characteristics | *M* | *SD* | Min | Max | $r_p$ | $p$ value |
| --- | --- | --- | --- | --- | --- | --- |
| Daily-life disturbance | 16.63 | 5.12 | 5 | 29 | .15 | .11 |
| Positive anticipation | 25.57 | 6.44 | 8 | 42 | -.02 | .80 |
| Withdrawal | 17.63 | 5.52 | 7 | 32 | -.06 | .54 |
| Cyber relation | 18.61 | 5.55 | 7 | 33 | -.04 | .69 |
| Overuse | 15.50 | 4.12 | 4 | 24 | .15 | .12 |
| Tolerance | 9.70 | 3.49 | 3 | 18 | -.01 | .90 |
| Total scores | 103.65 | 22.63 | 45 | 172 | .03 | .76 |

Participants were also asked to indicate phone conscious thought by responding to the following statement: “during the memory task, how often do you think of your smartphone”. Results showed a significant negative relationship between phone conscious thought and memory score, *r*_*S*_ = −.25, *n* = 119, *p* = .01. We anticipated a higher phone conscious thought for the LS group since their phone was kept away from them during the task and examined the relationship for each condition using a Spearman’s rho coefficient. Results showed a significant negative relationship between phone conscious thought and memory accuracy in the HS condition, *r*_*S*_ = −.49, *n* = 61, *p* = < .001, as well as the LS condition, *r*_*S*_ = −.27, *n* = 58, *p* = .04.

### Affect/Mood changes after being separated from their phone

We anticipated that our participants may have experienced either an increase in negative affect (NA) or a decrease in positive affect (PA) after being separated from their phone (LS condition). For LS participants, a paired sample t-test showed a significant decrease in participants’ positive affect before (*M* = 31.12, *SD* = 5.79) and after (*M* = 29.36, *SD* = 6.58) completing the memory task, *t* (57) = 2.48, *p* = .02, but not for the negative affect before (*M* = 20.71, *SD* = 6.49) and after (*M* = 21.19, *SD* = 6.84) completing the memory task, *t* (57) = .69, *p* = .50. A similar outcome is also shown in the HS condition, in which there is a significant decrease in positive affect only, *t* (60) = 3.45, *p* = .001 (see Fig 3).

**Fig 3.**
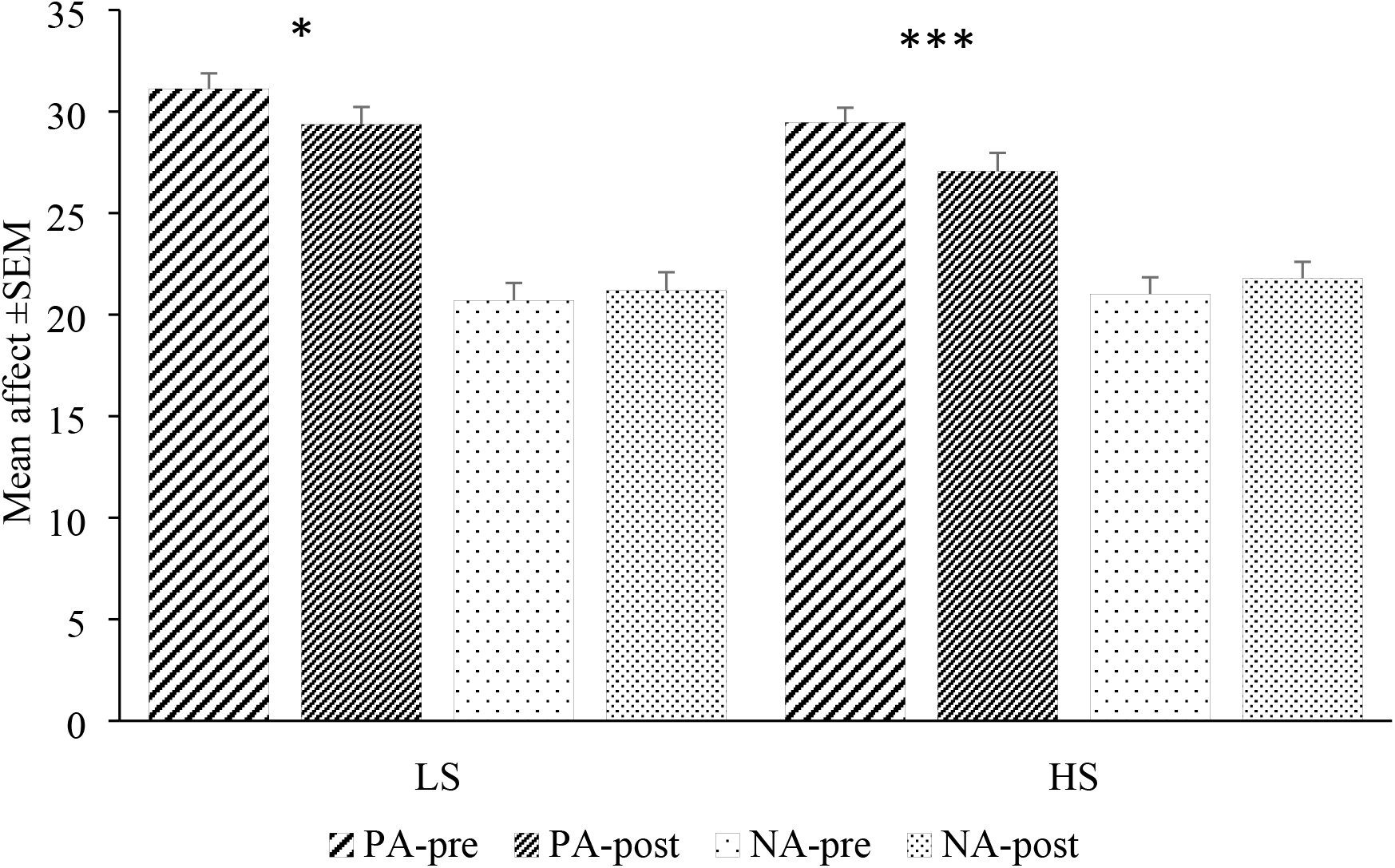
Mean positive affect (PA) and negative affect (NA) pre- and post-memory task between low phone salience (LS) and high phone salience (HS) groups (n = 119) *** p < .001, * p < .05.

### Relationship between phone conscious thought, smartphone addiction scale, and mood changes to memory recall accuracy

Preliminary analyses were conducted to ensure no violation of the assumptions of normality, linearity, multicollinearity and homoscedasticity. There was a significant positive relationship between SAS scores and phone conscious thought, *r*_*S*_ = .25, *n* = 119, *p* = .007. Using the enter method, we found that phone conscious thought explained by the model as a whole was 19.9%, *R*^2^ = .20, *R*^2^_*Adjusted*_ = .17, *F* (4, 114) = 7.10, *p* < .001. Phone conscious thought significantly predicted memory accuracy, *b* = −.63, *t* (114) = 4.76, *p* < .001, but not for the SAS score, *b* = .02, *t* (114) = 1.72, *p* = .09, PA difference score, *b* = .05, *t* (114) = 1.29, *p* = .20, and NA difference score, *b* = .06, *t* (114) = 1.61, *p* = .11.

### Perception between phone usage and learning

For the participants’ perception on their phone usage on their learning “In general, how much do you think your smartphone affects your learning performance and attention span?”, there was no significant difference between LS (*M* = 4.22, *SD* = 1.58) and HS participants (*M* = 4.07, *SD* = 1.62), *t* (117) = .54, *p* = .59. There was also no significant correlation between perceived cognitive interference and memory accuracy, *r* = .07, *p* = .47.

## Discussion

Our aim was to examine the effect of mobile phone presence on learning and memory, and in addition, to investigate the relationship between smartphone addiction, mood states and attachment to memory recall accuracy. First, our findings are consistent with prior studies (11–13) for participants had lower accuracy when their phone was next to them (HS), and higher when separated from their phones (LS). The mere presence of a phone even when not in use sufficiently distracted them in a simple learning and memory task.

Second, our findings showed that participants performed poorer in the memory task if they thought about their phone more often (high phone conscious thought) as phone conscious thought explained close to 20% of the variance in predicting memory accuracy scores. Our findings on phone conscious thought is consistent with Ward et al. (12) results, in that cognitive performance was moderated by high attachment to their phones. Although we did not find a significant relationship between SAS to memory accuracy, our measurement of ‘phone conscious thought’ is more relevant and meaningful because it measured participants’ separation anxiety from their phone at present time more directly, and higher scores on this item are also indicative of addiction (15). The similar high scores in the SAS for both HS and LS participants is a reflection of our sample population, and is similar to the ones reported in Kwon’s et al. (23) study, suggesting that addiction is relatively high in the student population compared to other categories such as employees, professionals, unemployed. Unlike other age groups, students use their cognitive resources more extensively to cope with life in general and is less able to manage their time effectively (24). The high SAS scores and primary use of phone for social media also indicates that the participants could be ‘problematic users’ as identified by Panek et al. (25). Frequent checks on social media is an indication of lower levels of self-control and having higher levels of need for belonging. Further, students’ usage of social networking (SNS) is a way of being common in them and the fear of missing out (FOMO) may fuel the SNS addiction (26). Our findings showed a decrease in PA after completing the tasks in our sample. Taken together with the high SAS and high phone conscious thought, the reduced PA in our participants is likely to have stemmed from the prohibited usage and/or separation from their phone. This is consistent with Cheever et al. (14) for their participants reported increased anxiety over time when separated from their phones. To sum it, Clayton, Leshner and Almond (17) proposed that people see mobile devices as an extended part of themselves, and that the inability to use their phone results in reduced sense of self, as if losing one’s limb.

### Further Studies

Future studies should look into the online learning environment. Students are often users of multiple devices and are expected to use their electronic devices frequently to learn various learning materials. Because students frequently use their mobile phones for social media and communication during lessons (25,27), the online learning environment becomes far more challenging compared to a face-to-face environment. It is highly unlikely that we can ban the phones despite evidence showing that students performed poorer academically with their phones next to them (7). The challenge is to engage students on their phones as a form of immersive learning experience, thus minimising other content while keeping students focused on their lessons. Some online platforms e.g. Kahoot and Mentimeter create a fun interactive experience to which students complete tasks on their phones and allow the instructor to monitor their performance from a computer. Another example is to use Twitter as a classroom tool (28).

Our findings report that the most frequently used feature is the SNS sites e.g. Instagram, Facebook, and Twitter. These behaviours are likely to remain the same when students graduate and move into the workforce. However, the work and study environments come with different set of norms and rules, and one study has shown that new entries into the workforce are unable to maintain proper barriers between social and professional lives (29) e.g. SNS as alternate academic platforms (30). Yet, SNS can be useful to boost productivity at work, but requires more monitoring (31) especially for our young workforce. That said, businesses and line managers need to reconsider the use of SNS that provides a good balance between social and professional demands for the young workforce.

## Conclusion

We cannot deny the ubiquitous nature of mobile phones in our lives, but the distraction seems to permeate in our daily tasks. Our findings support that the physical presence of mobile phones was distracting in a simple learning and memory task, and having no interaction on the phone for approximately 20 minutes reduced positive mood. Further, frequent thoughts about the phone is indicative of an attachment behaviour to the phone which contributed to overall poorer memory performance. With the rapid rise in e-learning environment and increasing phone ownership, mobile phones will continue to be present in the classroom and work environment. It is important that we manage or integrate the phones into the classroom but the extent of the device purpose will remain as a contentious issue between instructors and students.

## Acknowledgements

We would like to thank our participants for volunteering to participate in this study.

